# Multi-Tissue Transcriptome-Wide Association Study Identifies 26 Novel Candidate Susceptibility Genes for High Grade Serous Epithelial Ovarian Cancer

**DOI:** 10.1101/330613

**Authors:** Alexander Gusev, Kate Lawrenson, Felipe Segato, Marcos A.S. Fonseca, Siddhartha Kar, Kevin C. Vavra, Janet M Lee, Tanya Pejovic, Ovarian Cancer Association Consortium, Beth Y. Karlan, Matthew L. Freedman, Houtan Noushmehr, Paul D.P. Pharoah, Bogdan Pasaniuc, Simon A. Gayther

## Abstract

Genome-wide association studies (GWASs) have identified about 30 different susceptibility loci associated with high grade serous ovarian cancer (HGSOC) risk. We sought to identify potential susceptibility genes by integrating the risk variants at these regions with genetic variants impacting gene expression and splicing of nearby genes. We compiled gene expression and genotyping data from 2,169 samples for 6 different HGSOC-relevant tissue types. We integrated these data with GWAS data from 13,037 HGSOC cases and 40,941 controls, and performed a transcriptome-wide association study (TWAS) across >70,000 significantly heritable gene/exon features. We identified 24 transcriptome-wide significant associations for 14 unique genes, plus 90 significant exon-level associations in 20 unique genes. We implicated multiple novel genes at risk loci, e.g. *LRRC46* at 19q21.32 (TWAS *P*=1×10^−9^) and a *PRC1* splicing event (TWAS *P*=9×10^−8^) which was splice-variant specific and exhibited no eQTL signal. Functional analyses in HGSOC cell lines found evidence of essentiality for *GOSR2, INTS1, KANSL1* and *PRC1*; with the latter gene showing levels of essentiality comparable to that of *MYC*. Overall, gene expression and splicing events explained 41% of SNP-heritability for HGSOC (s.e. 11%, *P*=2.5×10^−4^), implicated at least one target gene for 6/13 distinct genome-wide significant regions and revealed 2 known and 26 novel candidate susceptibility genes for HGSOC.

**STATEMENT OF SIGNIFICANCE:** For many ovarian cancer risk regions, the target genes regulated by germline genetic variants are unknown. Using expression data from >2,100 individuals, this study identified novel associations of genes and splicing variants with ovarian cancer risk; with transcriptional variation now explaining over one-third of the SNP-heritability for this disease.

## INTRODUCTION

Invasive epithelial ovarian cancer (EOC) is a heterogeneous disease with a major heritable component (1). There are several histological subtypes of invasive EOC, each associated with different genetic and epidemiological risk factors, clinical features and likely cells of origin; high grade serous (HGSOC) is the most common histotype, representing about two-thirds of cases. Highly penetrant germline mutations in the homology directed repair (HDR) genes including *BRCA1* and *BRCA2* are the most significant genetic risk factors for HGSOC, and are often associated with familial clustering of EOC cases (1). However, mutations in these genes are rare in most populations, and only account for about 10% of cases. In total, the known susceptibility genes account for less than 40% of the heritable risk of EOC. A substantial proportion of the remaining EOC risk is likely to be due to alleles of greater frequency in the population but of lower penetrance. Over the last few years, genome-wide association studies (GWAS) have identified 39 different regions of the genome associated with EOC risk, mainly in European populations; but evidence from GWAS for other phenotypes suggest that hundreds of additional common low penetrance EOC risk alleles are likely to exist and await identification (2‐4).

Typically, SNPs associated with disease risk are located in the non-protein coding genome suggesting they function through interactions with non-coding elements (e.g. epigenomic modifications, non-coding RNAs) that regulate gene expression. However, because there is no precise genetic code for the non-protein coding genome, there are neither reliable *in silico* algorithms to predict the functional effects of risk associated SNPs, nor their likely target genes. Several studies have indicated that the target genes at risk loci are often in close proximity to the causal risk-associated SNP (i.e. they are *cis*-regulated)(5‐8). Expression quantitative trait locus (eQTL) analysis can be used to interrogate the associations between risk genotypes and gene expression and several studies have successfully used this approach to indicate the likely susceptibility gene at known GWAS risk loci (5‐8). These studies have identified several novel candidate EOC susceptibility genes in risk regions identified by GWAS (5‐12); but unlike high penetrance HGSOC susceptibility genes that function in HDR pathways, the genes associated with common, non-coding EOC risk variants (e.g. *CDC42, CDCA8, PAx8, HOxD9, CHMP4C, OBFC1, RCCD1* and *ABHD8*) more often function in cell cycle control pathways or gene regulation and not DNA repair.

More recently, transcriptome-wide association approaches (TWAS) have been developed that leverage expression data by ‘imputing’ gene expression across a large cohort of genotyped individuals to identify target genes associated with phenotypes of interest (13‐17). For a given gene, TWAS is a test of local genetic association between gene expression and GWAS risk. Under a model where genetic variation alters gene expression and leads to changes in disease risk, TWAS will identify genes whose genetically regulated expression is associated to disease risk. TWAS may additionally increase power versus single SNP association testing either by reducing the multiple testing burden or aggregating multiple expression-altering variants into a single test. However, TWAS may also be significant due to pleiotropy between the expression-altering and risk-altering variants or variants they tag. TWAS is therefore a first step to prioritize among putative target genes, with experimental validation needed to establish causality.

In the current study we established the most comprehensive genome-wide genotype-gene expression datasets available, with >2,000 samples including primary HGSOCs, EOC precursor tissues (ovarian and fallopian epithelial cells) and other hormonal-related cancers (breast and prostate cancer), to perform TWAS analysis to specifically identify candidate susceptibility genes associated with EOC risk alleles. Finally, functional information from a gene knockout screen (18) was used to identify candidate susceptibility genes with a functional role in HGSOC cells.

## RESULTS

### Genetic control of gene expression is reprogrammed during tumorigenesis

We first sought to investigate the genetic control of gene expression in EOC precursor tissues and HGSOCs. We assayed genotype, gene expression, and quantified splicing data for 115 primary OSECs and 70 primary FTSECs, as well as 394 HGSOCs from The Cancer Genome Atlas (TCGA). The majority of HGSOCs are thought to derive from fallopian tube secretory epithelial cell (FTSEC) precursors (19‐23), although a subset may also arise from ovarian surface epithelial cells (OSECs) (24‐29). We first quantified the SNP-heritability (h^2^_*g*_) and genetic correlation (*r*_*g*_) of gene expression and splicing between pairs of tissues (see Methods). For a given tissue, cis-*h*^2^_*g*_ is defined as the fraction of phenotypic variance explained by SNPs within 500 kb of the gene boundary. For a pair of tissues, cis-*r*_*g*_ is defined as the correlation of causal genetic effects on expression across all SNPs within 500 kb of the gene boundary. The average cis-(*h*^2^_*g*_) was significant in all tissues with an average of 0.026 for overall expression and 0.016 for splice variation, as has been generally observed across diverse tissues (**Table S1**). Mean cis-*r*_*g*_ was highest between FTSECs and HGSOCs (*r*_*g*_ = 0.071, standard error [s.e.] = 0.031) compared to OSECs and HGSOCs (*r*_*g*_ = −0.022, s.e. = 0.029). We observed a similar, albeit non-significant, trend for heritable splicing events, with a genetic correlation of 0.024 (s.e. = 0.016) between HGSOCs and FTSECs, and −0.018 (s.e. = 0.013) between OSECs and HGSOC. These results are consistent with FTSECs representing the predominant cell-of-origin for HGSOC. Genetic correlation of both overall gene expression and splicing events was substantially higher when normal OSECs and FTSECs were compared (*r*_*g*_= 0.359, s.e. = 0.046 for overall expression; *r*_*g*_ = 0.302, s.e. = 0.023 for splicing) indicating that genetic control of gene expression is altered during tumorigenesis. Lastly, we evaluated *r*_*g*_ between four molecular subtypes of HGSOCs characterized by gene expression signatures by TCGA (30) but observed no significant difference from 1.0 (**Table S2**). We therefore treated all HGSOCs as a single group for all subsequent analyses.

### Overview of TWAS and statistical methods

Next, we performed a transcriptome-wide association study (TWAS) using diverse gene expression models and GWAS statistics from 13,037 HGSOC cases and 40,941 controls estimated by the Ovarian Cancer Association Consortium (OCAC). We first constructed the predictive models of gene expression and splicing using genotypes and expression from the EOC precursor and HGSOC expression data described above, as well as tumor datasets for other hormonally-regulated cancers (breast and prostate cancers) that have recently been shown known to exhibit genetic pleiotropy with EOC: Breast tumors (n = 1,027) with matched normal precursor tissues (n = 80); and prostate tumors (n = 483) (5,12,31). All data sets were assayed by RNA-seq. We additionally evaluated exon-exon junction usage as a measure of alternative splicing (see Methods). For each gene/exon event in a given tissue, we estimated cis-*h*^*2*^_*g*_ using a variance-component model from all common variants within the *cis h*^*2*^_*g*_ locus and retained all gene-tissue combinations that were nominally heritable (cis-*P* <0.01). A total of 12,514 significantly heritable genes and 55,618 significantly heritable exon events were identified across all cohorts. We then trained multiple predictive models using all SNPs in the locus, including ridge regression / BLUP, lasso, elastic net, and the Bayesian Sparse Linear Mixed Model (BSLMM). For each model, predictive accuracy was evaluated by five-fold cross-validation against the actual measured expression. Mean cross-validated predictor R^2^ was 0.062 and highly significant (median *P* = 1.8×10^−4^ across all predictors), consistent with previous findings that low average heritability of gene expression can be compensated for by sample size sufficient to produce reliable genetic predictors (**Table S3**). For genes that were significantly heritable in multiple tissues, we evaluated their concordance by predicting into a held-out reference cohort of 1000 Genomes Europeans. These predictors were significantly correlated between both tumor and normal tissue pairs (mean correlation across all tissue pairs = 0.554, s.e. = 0.0057), underscoring their reproducibility in independent data (**Table S4**).

We then tested each gene/exon prediction model for association with risk using HGSOC GWAS summary data. The genetically predicted expression of 24 gene-level models and 90 exon-level models were significantly associated with risk after Bonferroni correction for 68,132 total tests (**Supplementary Fig. S1**). We describe detailed analysis of these associations below. TWAS may identify co-incidental genetic associations due to partial tagging between the expression and disease causing variants, and so we performed additional statistical analyses on a locus-by-locus basis to assess this possibility. First, we condition every GWAS association on the predicted value of each significant TWAS gene to assess how much association signal remains independent of the TWAS association (see Methods). Residual significant signal after conditioning is an indicator that the TWAS association is partially tagging the causal variants, or that other independent causal variants are present at the locus (as with SNP-based conditioning). Second, we perform a “colocalization” analysis using the COLOC software (58), which evaluates the posterior probability that the genetic association to the gene/exon is driven by a single shared causal variant with the GWAS risk association (termed “PP4” in the COLOC notation). This model does not consider colocalization between multiple causal variants, so high PP4 is a more stringent threshold to clear than the TWAS association and may miss true colocalization at loci with heterogeneous effects on expression and disease. COLOC additionally estimates the probability that the expression and GWAS are driven by two distinct causal variants (PP3), and we use low PP3 as a less stringent threshold for evidence of non-independent association signal (though it may still be confounded by multiple causal variants).

### Gene-level associations validate previous eQTLs and identify novel susceptibility genes for HGSOC

First we characterized gene level events, identifying 24 TWAS associations (14 unique genes) across the six tissue types after Bonferroni correction (**Table 2, Supplementary Table S5, Fig. S1**). The number of significant associations was strongly correlated with the number of tested genes (R^2^=0.56) suggesting that expression reference size and quality, rather than tissue specificity, is the main factor driving TWAS gene discovery (**Fig. S2**). A single association was detected in FTSECs: expression of *LRRC46* at 17q21.32 (TWAS *P* = 1.5×10^−9^). Four genes were associated with risk in HGSOCs, all located within an inversion at chromosome 17q21.31 - *KANSL1, LRRC37A2, ARL17A* and *RP11-259G18.1 (32)*. *ARL17A* was a notable example where ovary-specific eQTLs explained the local GWAS signal, but significant eQTLs observed in breast and prostate were independent (**Fig. 1**). *ARL17A* has not been previously implicated in ovarian cancer, although *KANSL1- ARL17A* gene fusions have been implicated in pancreatic cancer (33). Two previously reported eQTLs at EOC risk loci identified in ovarian tissues were also detected in other tissue types - *CHMP4C* at the 8q21 locus (9) was detected in breast and prostate tumors (TWAS *P* = 1.3×10^−10^ and 9.8×10^−10^, respectively) and *ANKLE1* at the 19p13 locus (5) was detected in prostate tumors (TWAS *P* = 1.6×10^−13^). A follow-up colocalization analysis showed that 12/24 TWAS associations exhibited strong evidence of a single shared causal variant (PP4 > 0.8) and only 5/24 had evidence of joint causal variants (PP3 > 0.2).

**Table 1.** Genetic correlation of expression and splicing in ovarian tissues. Genetic variance/covariance was estimated across all significantly heritable genes (in any panel) using HE-regression, averaged, and transformed to genetic correlation (*r*_*g*_). Standard error shown in parentheses.

| Expression | | Overall $r_g$ (se) | Splicing $r_g$ (se) |
| --- | --- | --- | --- |
| Tumor HGSOC | Normal OSEC | -0.022 (0.029) | -0.018 (0.013) |
| Tumor HGSOC | Normal FTSEC | 0.071 (0.031) | 0.024 (0.016) |
| Normal OSEC | Normal FTSEC | 0.359 (0.046) | 0.302 (0.023) |

**Table 2.** TWAS analyses in ovarian cancer. FTSEC, fallopian tube secretory epithelial cell; OSEC, ovarian surface epithelial cell; HGSOC, high-grade serous ovarian cancer.

| Site | Type | Samples | Tested genes | Tested splice variants | Significant genes | Significant splice variants | Significant splice variant genes |
| --- | --- | --- | --- | --- | --- | --- | --- |
| Breast | Normal tissue | 80 | 769 | 3485 | 3 | 21 | 4 |
| Breast | Tumor tissue | 1027 | 4215 | 21582 | 6 | 40 | 12 |
| Ovary | Normal FTSEC | 70 | 572 | 1409 | 1 | 0 | 0 |
| Ovary | Normal OSEC | 115 | 541 | 1399 | 0 | 0 | 0 |
| Ovary | Tumor tissue (HGSOC) | 394 | 2956 | 12580 | 4 | 9 | 6 |
| Prostate | Tumor tissue | 483 | 3461 | 15163 | 10 | 20 | 8 |

**Figure 1.**
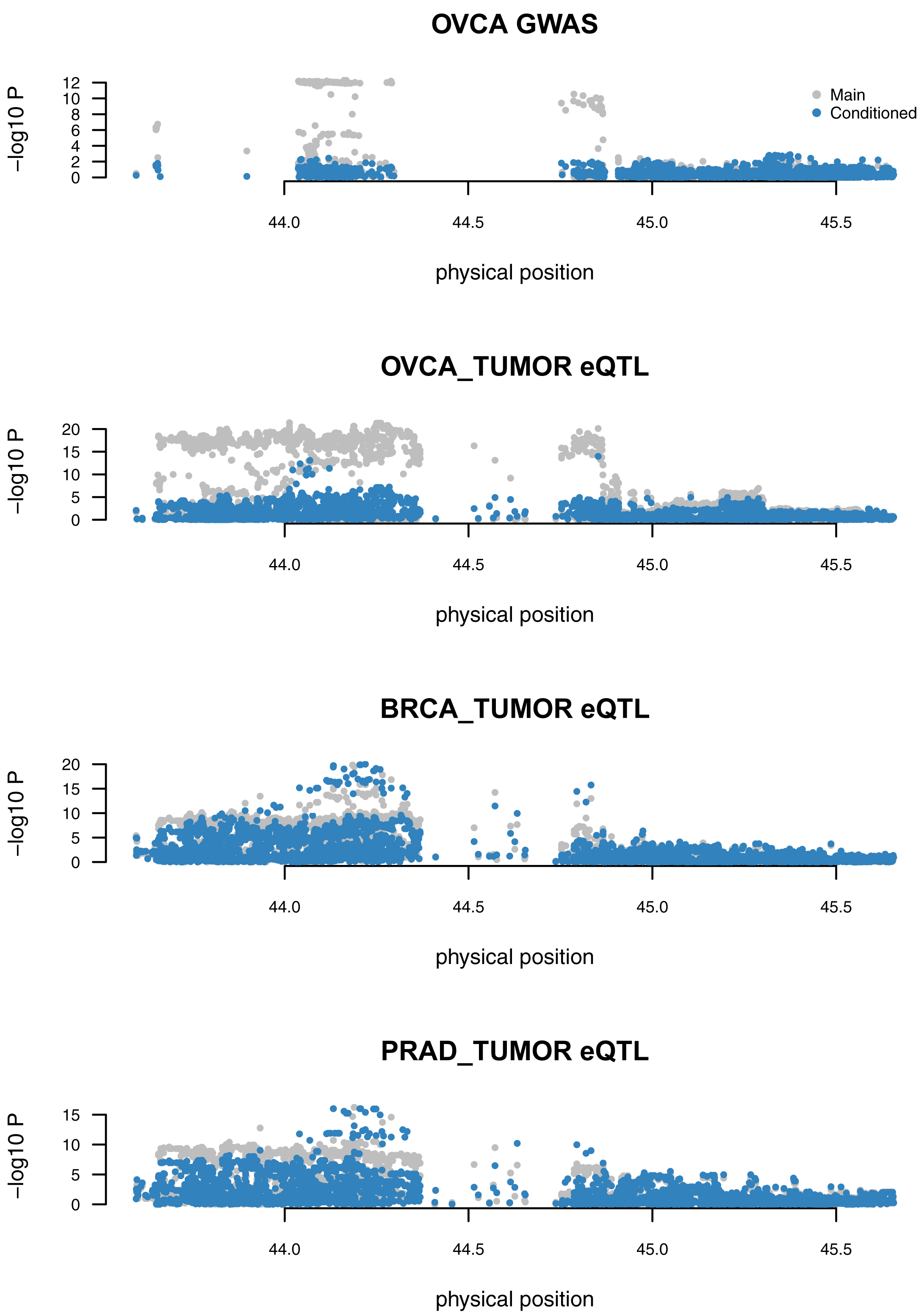
Ovary-specific TWAS association for *ARL17A*. The *ARL17A* gene is under strong genetic control in multiple tissues but only colocalizes with GWAS in ovarian tumors. Each panel shows Manhattan plot before (gray) and after (blue) conditioning on the TWAS predictor trained on ovarian tumor expression: **A,** GWAS associations, with signal fully explained after conditioning on the predictor; **B,** ovarian tumor eQTLs; **C,** breast tumor eQTL; **D,** prostate tumor eQTL. **A** and **B** show associations are explained by the ovarian TWAS predictor, whereas **C** and **D** show associations are independent of the ovarian TWAS predictor.

We identified two genes that were more significantly associated than any single overlapping SNP: *KANSL1* and *RP11-259G18.1* at the 17q21.31 locus, both in ovarian tumors. The increased significance is indicative of multiple eQTL signals aggregating risk heritability to increase significance. In particular, the *KANSL1* association, driven by 4 SNPs in an elastic net predictive model, was substantially higher than the best overlapping SNP: TWAS *P* = 6.9×10^−11^ *versus* GWAS *P* = 8.9×10^−9^. The multi-SNP *KANSL1* predictive model had a cross-validated R^2^ with measured expression of 0.65 (*P* = 6.5×10^−85^), compared to the top eQTL having an R^2^ of 0.49 (*P* = 1.4×10^−40^), indicative of significant secondary effects on expression and further supporting a multi-eQTL/GWAS hypothesis. Conditioning each SNP in the broader locus on the predicted expression of *KANSL1* reduced the lead GWAS SNP association from *P* = 1.1×10^−11^ to *P* = 0.04 and accounted for all genome-wide significant signal.

We replicated nine out of the 14 TWAS associations using predictors from the Genotype-Tissue Expression (GTEx) study (after Bonferroni correction for 84,964 GTEx predictors tested; **Supplementary Table S6**). Only two were significant in whole ovary tissues analyzed in GTEx - the paralogs *LRRC37A* and *LRRC37A2*, which were significant in nearly all tissues except for testis and normal prostate tissues. Notably, *ARL17A*, while highly significant in HGSOCs was not significant in most GTEx tissues, suggesting tumor-specific regulatory effects. The GTEx analysis identified two additional transcriptome-wide significant genes that did not lie in genome-wide significant HGSOC loci. First, *RCCD1* at 15q26 was significant in 20 tissues (including breast); this locus had previously been identified in a meta-analysis of breast and ovarian cancer and this gene reported as the putative target based on GTEx eQTL analyses (12). Second, *DNALI1* at 1p34 was significant in 9 tissues (but not in breast, prostate or ovary); this locus was previously reported as genome-wide significant in an analysis of serous EOC where the *RSPO1* gene was identified as a putative target but did not show an eQTL association (34). At both loci, conditioning on the predicted expression accounted for all genome-wide significant signal, consistent with these genes being potential mediators of the association (**Supplementary Fig. S3, S4**).

### Multi-tissue exon-level associations implicate novel transcripts in HGSOC development

Next, we performed a splice-TWAS across all significantly heritable exon-exon usage features, identifying 90 splice-TWAS associations with EOC risk in 20 unique genes (after Bonferroni correction; **Table 2, Supplementary Table S7**). This included 14 genes that did not have a significant gene-level TWAS association the TWAS analysis of overall expression. The mean splice-TWAS effect-size was significantly negative (Z-score of −4.2 s.e. 0.50) suggesting that variants decreasing exon usage are more typically associated with increased risk. Co-localization analysis (58) showed that 67/90 associations were consistent with a shared causal variant (posterior on shared > 0.8) and 84/90 were inconsistent with a single distinct causal variant (posterior on distinct <0.2). Five loci contained only a single significantly associated gene and we investigated these loci in detail.

In breast tumors we identified a splice-TWAS association for *PRC1* which fully explains the GWAS signal at the 15q26.1 locus (TWAS *P* = 8.9×10^−8^, PP4 = 1.0), a pleiotropic risk locus previously associated with both breast and ovarian cancer risk (12) (**Fig. 2**). Notably, no significant eQTLs were observed for overall expression of *PRC1*, highlighting a genetic effect on splicing that is independent of total expression. Predictors for total *PRC1* expression could be computed in three GTEx tissues, but none resulted in significant TWAS associations. We separately identified a significant TWAS association for the nearby *RCCD1* gene using expression from GTEx(see above), reproducing the previously identified putative target gene (12).

**Figure 2.**
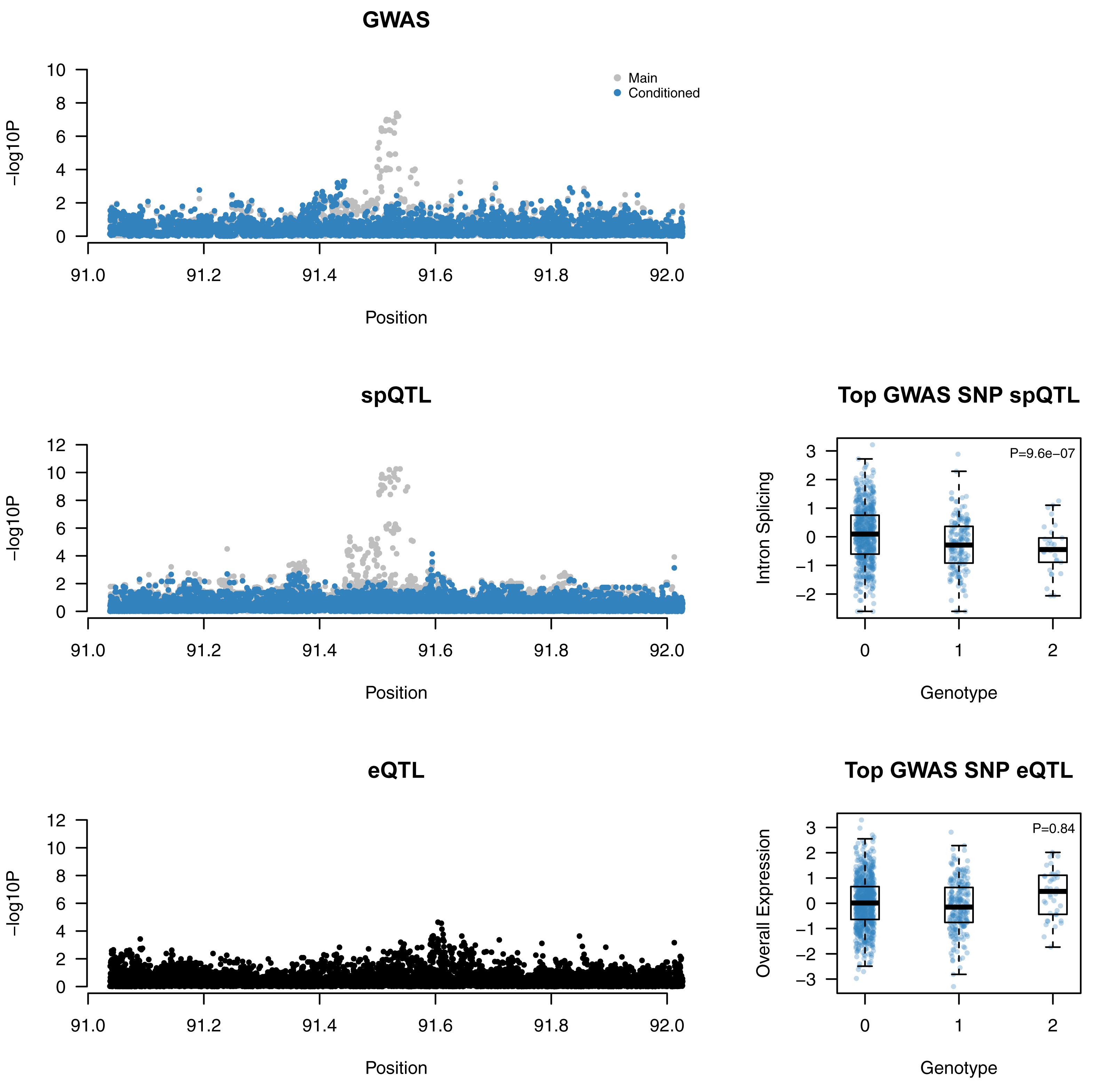
Splice-TWAS association at *PRC1* implicates novel target gene independent of genetic effects on total expression. Panels **A, B, D** show Manhattan plot before (gray) and after (blue) conditioning on the top splice-QTL. Panels **C, E** show box and scatter plots of normalized intron (top) and overall (bottom) expression, stratified by lead GWAS SNP genotype.

We detected four splice-TWAS associations for *CHMP4C*, a previously described eQTL in ovarian tumors (9); we now detected the same association in prostate tumors. The lead spQTL (rs74758321) is within 300bp of the splice junction and in perfect linkage disequilibrium with the top GWAS, making it a plausible causal candidate SNP in addition to the previously implicated missense eQTL.

Lastly, three loci with single gene associations had partial evidence that fully explained the locus. A significant splicing event was detected for *LEKR1* in HGSOCs (TWAS *P* = 1.5 ×; 10^−15^, PP4 = 0.96), which was the strongest splice-TWAS association with evidence of co-localization in our study. The lead spQTL for *LEKR1* (rs344076) in GTEx is also an eQTL for *LEKR1* in 20 tissues and *TiPARP* in 2 tissues. However, conditioning on the LEKR1 association did not account for all signal at this locus, indicating only partial tagging of the underlying causal variant. Likewise, splicing for *HOxD-AS1* in ovarian tumors was the only association at 2q31 but had moderate evidence of co-localization (PP4 = 0.78) and did not account for all signal in a conditional analysis. Besides *PRC1*, we observed one additional association that did not overlap a genome-wide significant region: splicing for *INTS1* in breast tumors at the 7p22 locus. However, the lead spQTL was only marginally significant (*P* = 1.2×10^−4^) and could not be confirmed by colocalization analysis (**Supplementary Table S7**, **Supplementary Fig. S5**). We therefore cannot consider this a true novel association without replication in an independent GWAS cohort.

The majority of associations (64/90) were at the 17q21.31 locus with a inversion polymorphism spanning˜900kb. This region contains hundreds of variants in high linkage disequilibrium that all represent putative causal alleles and are involved in genetic co-regulation of 14 genes, with evidence of multiple clusters of independent associations (**Supplementary Fig. S6A**). We observed a complex co-regulation of >3 unique genes at one other locus - 19p13.11 - with evidence of multiple independent associations. We performed stepwise conditional analysis of all significant TWAS/spTWAS associations in the locus to identify the minimal set of genes that jointly explained all of the genome-wide significant signal. The final model contained four jointly significant splicing events - *CPAMD8, MAP1S, OCEL1* - reducing the lead GWAS SNP from *P* = 7.8×10^−^25 to *P* = 1.4×10^−6^ (**Supplementary Fig. S6B, S7A**). The implication of multiple joint predictors suggests either the presence of multiple causal variants at this locus, or colocalization with a single gene that was not tested here. A splicing event for the *BABAM1* gene had the highest colocalization posterior in the locus (PP4 = 0.95) but did not account for all genome-wide significant association in a conditional analysis (**Supplementary Fig. S7B**). The 19p13.11 risk locus is also associated with triple-negative breast cancer (5) and *BABAM1*, a known *BRCA1*-interacting protein, is therefore a compelling target. Our previous gene-level functional studies failed to find compelling evidence to support a role for *BABAM1* in early ovarian and breast tumorigenesis and putatively implicated neighboring genes *ABHD8* and *ANKLE1* (5); our new analyses suggest characterization of the function of specific *BABAM1* splice variants is also warranted. In addition, further studies are needed to understand the apparently contradictory results from the conditional and colocalization analyses, suggestive of multiple causal variants or complex local haplotype structure. The extensive, cross-tissue genetic co-regulation underscores the complexity of these loci and motivates continued experimental fine-mapping.

Eight out of the 20 unique genes with a splice-TWAS association replicated within one of the other five tested tissues (after Bonferroni correction for 20×5 tests) (**Supplementary Table S8**). In particular, splicing for *CPAMD8, INTS1, SNx11, GOSR2, NFE2L1,* and *SP2* was heritable but not TWAS significant in at least one other tissue, indicative of tissue-specific regulation (rather than insufficient power to detect the QTL in the other tissue). Overall, the GWAS contained 13 contiguous genome-wide significant regions, of which 6 were within 500 kb of a TWAS or splice-TWAS association (**Supplementary Table S9**). These 6 regions implicated a total of 74 associated features out of a 1,156 tested, demonstrating a substantial number of heritable gene/tissue combinations that have also been ruled out as likely cis targets.

### Pleiotropic associations with breast cancer risk GWAS

We further tested the 114 transcriptome-wide significant features (90 splice events and 24 genes) for pleiotropic associations with breast cancer risk from a recent GWAS (3). 80 out of 114 features showed evidence of significant TWAS association (P<0.05/114), which indicates extensive pleiotropy between the breast and ovarian cancer at these loci. Of these, five were genome-wide significant (*P* <5×10^−8^) for breast cancer, all of which were exon-level events in breast tumors: one for *PRC1* and four for *LRRC37A2*. These results highlight two robust genome-wide significant loci associated with breast and ovarian cancer that also exhibit effects on splicing of the same genes. We repeated the same analysis for a recent GWAS for prostate cancer but did not identify any features with *P* <0.05, suggesting that the extensive expression-based pleiotropy we observe between breast and ovarian cancer is not expected by chance.

### Ovarian tumor gene expression and splicing explains the greatest portion of EOC SNP-heritability

We quantified the portion of GWAS heritability explained by all TWAS/splice-TWAS predictors in each tissue type. As with analyses of polygenic GWAS heritability (35), this approach does not restrict to transcriptome-wide significant associations but estimates the overall proportion of heritability that can be accounted for by all TWAS predictors. Ovarian gene expression and splicing explained 41% (*P* = 2.5×10^−4^) of EOC SNP-heritability, which is greater than for either breast or prostate cancer despite the smaller sample size of the EOC study (**Table 3**). We did not have sufficient power to distinguish the portion of expression heritability between ovarian tumor and normal tissues. We caution that this estimate includes any tagged genetic effects that alter expression and risk independently, and should thus be interpreted as an upper bound.

**Table 3.** Ovarian cancer heritability explained by *cis*-regulated gene expression from combined tumor/normal predictors.

| | $h^2_g$ | se | p-value | % $h^2_g$ | # features | $h^2_g$ per feature |
| --- | --- | --- | --- | --- | --- | --- |
| Ovarian | 0.015 | 0.0041 | $2.5 \times 10^{-4}$ | 41% | 19,501 | 7.7E-07 |
| Prostate | 0.009 | 0.0033 | $6.5 \times 10^{-3}$ | 25% | 18,646 | 4.8E-07 |
| Breast | 0.014 | 0.0041 | $6.5 \times 10^{-4}$ | 38% | 29,326 | 4.8E-07 |

### Functional characterization of candidate susceptibility genes

Finally we explored the functional role of the 28 candidate susceptibility genes identified through our TWAS and spTWAS analyses. We mined publicly available data from a gene essentiality screen (18); knockout data were available for 24 genes in 13 HGSOC cell lines (**Figure 3**). Four genes showed evidence of essentiality (CERES Score <0.5) - golgi SNAP receptor complex member 2 (*GOSR2,* median CERES score = −0.72, s.d.= 0.11), integrator complex subunit 1 (*INTS1*, median CERES score = −0.97, s.d. = 0.10), KAT8 regulatory NSL complex subunit 1 (*KANSL1*, median CERES score = −0.53, s.d. = 0.15) and protein regulator of cytokinesis 1 (*PRC1*, median CERES score = −1.17, s.d. = 0.14), with the latter gene showing similar levels of essentiality as *MYC*, a key oncogenic transcription factor in many tumor types, including ovarian (36). The mean CERES score across significant splice-TWAS genes (−0.24 s.e. 0.02) was significantly lower (more essential) than that of significant TWAS genes not associated through splicing (−0.08 s.e. 0.01; P=4.4×10^−5^ for difference by Wilcoxon rank sum test), suggesting that risk variants that affect splicing (and thus actual protein structure) may be more relevant to OvCa cell essentiality than risk variants that affect transcription (i.e. protein abundance). We caution, however, that the CERES data may not be reflecting changes to the precise transcript that is implicated by the splice-TWAS.

**Figure 3.**
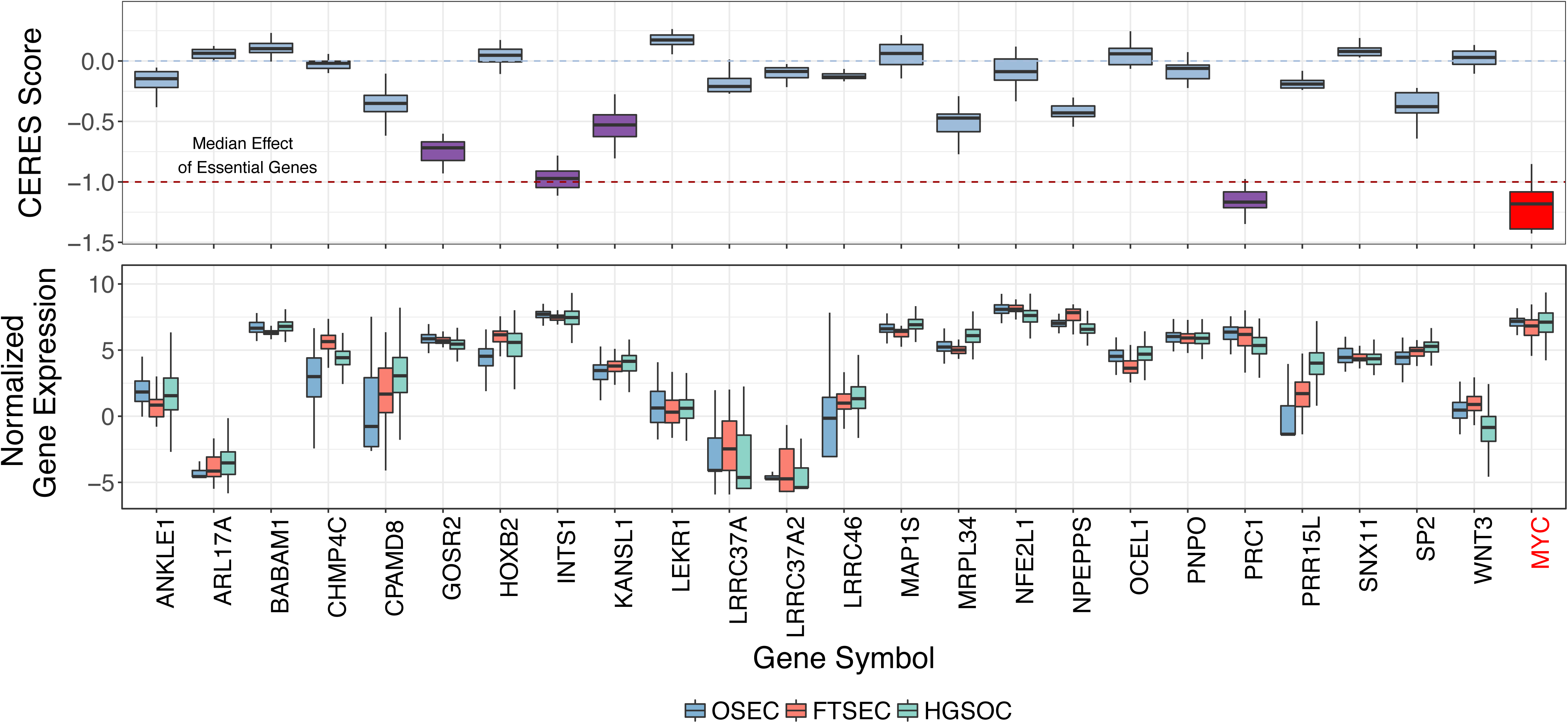
Functional analyses show evidence of essentiality for 3 TWAS/spTWAS genes. **A,** Gene knockout experiments in 13 HGSOC cell lines to determine gene essentiality. CERES Score is a copy number corrected indicator of depletion of gene-targeting guide RNAs, the lower the CERES Score, the more essential the gene. *MYC* is a known essential gene and is included as a positive control. CERES Score thresholds corresponding to the median score for non-essential and essential genes are indicated with a blue dashed line at 0 and red dashed line at −1, respectively. Genes with a CERES Score ≤ −0.5 (and therefore showing evidence of essentiality) are highlighted with purple boxes. **B,** Relative expression of each gene in OSECs, FTSECs and HGSOCs. In each plot the center line denotes the median, box limits represent upper and lower quartiles, whiskers show 1.5x interquartile range and any discrete data points show outliers.

## DISCUSSION

Genome-wide association studies (GWAS) have identified multiple of common variants associated with risks of high grade serous ovarian cancer (HGSOC); but for the vast majority of risk variants the underlying functional mechanisms, including the susceptibility genes targets, are unknown. In this work we integrate tissue specific gene expression and genotyping data with the largest GWAS available for HGSOC to identify susceptibility genes for HGSOC. We functionally validate 24 candidate susceptibility genes and find four genes to show evidence for essentiality.

Key to our approach is transcriptome measurements in tissues that are relevant to HGSOC pathogenesis; and so we compiled whole genome genotyping and gene expression data from 2,169 different patients representing 6 different tissue types. This represents the largest and most comprehensive gene expression and genotyping catalogue so far compiled to identify genotype-gene expression associations for ovarian cancer, and includes previously unpublished gene expression profiling data for normal precursor tissues for HGSOC, and primary ovarian, breast and prostate tumors. Transcript expression can be highly tissue and disease specific, and genotype-gene expression associations are also likely to be tissue specific. While our transcriptome-wide significant findings were primarily driven by reference panel size, our polygenic findings support this: a greater proportion of GWAS heritability was explained by all expression in the ovarian tumor tissues rather than breast and prostate tumors. Moreover, the genetic correlation between FTSECs and HGSOCs was higher than between OSECs and HGSOCs; published data strongly suggest that FTSECs are the most common precursor cell type for HGSOC (19‐23), although some data suggest that OSECs may be a precursor cell type for specific molecular subgroups of HGSOC (37‐39). Furthermore, only 2/21 genes we identified were transcriptome-wide significant in ovary tissues from the GTEx Consortium (with 11/21 significant across all GTEx tissues); likely because whole ovaries are predominantly composed of stromal (rather than epithelial) cells and the sample size in GTEx is small (n=54).

We conclude with several caveats of our approach. It remains likely that our TWAS analysis missed a unquantifiable proportion of true associations, while some associations may represent false positive findings due to chance co-regulation. HGSOCs are genomically unstable and accumulate copy number variations during their development, such variation will likely influence the patterns of gene expression and consequently the outcome of our TWAS analyses. Also, in building catalogues of gene expression profiles for multiple tissue types, we may not have included all relevant tissues, such as microenvironmental cell populations that influence disease risk *via* non-cell autonomous pathways (e.g. immune cells). Finally, compared to other TWAS studies, our study was limited by sample size and therefore statistical power to identify genotype-gene expression associations. Future studies to improve the power of TWAS analysis in ovarian cancer will need to establish substantially larger gene expression and genotyped datasets for normal precursor tissues, for HGSOC and for other EOC histotypes that were not evaluated in the current study.

Pleiotropy, defined here as a risk locus that shows evidence of susceptibility to two or more phenotypes, has emerged as a feature of GWAS risk loci. We have recently performed meta-analyses of GWAS datasets for different phenotypes to identify loci associated with risks of ovarian, breast and prostate cancer (12). This suggests that the functional mechanisms (and target susceptibility genes) underlying pleiotropic risk loci are likely to be similar for the different phenotypes. Thus, incorporating transcriptomic datasets for the ovarian, breast and prostate tumors in our TWAS may identify susceptibility genes that are associated with the different cancers. Consistent with this hypothesis, we observed a striking overlap between significant TWAS genes in GWAS for HGSOC and a recent GWAS of breast cancer (3). These findings merit further studies to identify evidence of pleiotropy and common cancer susceptibility genes for these two cancers.

Most risk variants identified by GWAS lie in non-protein coding regions and are postulated to influence the expression of target susceptibility genes *in cis*. Variants can alter gene expression through multiple mechanisms. Some SNPs may modify levels of primary transcript expression or structure (e.g. genetic variation in gene promoters, distal regulatory elements and splicing), while other variants may modulate post-transcriptional control of gene expression (e.g. variants in long non-coding RNAs or microRNAs). In the current analysis, we performed a TWAS to evaluate associations between genetic variation and both primary transcript expression (eQTL) and splice site variation (spQTL). Our TWAS validates previously described eQTL associations, including *CHMP4C, ANKLE1, and RCCD1 (5,9)*, and also identified candidate genes at risk loci where previous *cis*-eQTL studies have been unsuccessful, such as *LEKR1* splicing at the 3q25 locus. Moreover, the splice-TWAS analysis identified 19 genes that were not implicated by overall TWAS, nearly doubling the number of susceptibility genes. This included *PRC1* (at 15q26.1) which explained all of the GWAS signal while exhibiting no eQTL association and was not previously identified in an eQTL-based analysis. This highlights the power of TWAS to identify novel candidate susceptibility genes. Our findings indicate that primary transcript expression is not the sole mechanism by which target gene expression can be altered at GWAS susceptibility loci, and that QTL-based studies need to evaluate alternative mechanisms to find the target genes associated with common risk variants.

Establishing a functional role for candidate genes and their associated risk alleles in disease is beyond the scope of the current study. However for each candidate gene we evaluated the evidence for their possible role in cancer biology from their known or predicted functions. High penetrance susceptibility genes so far identified in the development of HGSOC - *BRCA1, BRCA2, BRIP1, RAD51C, RAD51D* (1) - are all involved in double-strand DNA break repair. However, for EOC, the genes associated with low penetrance risk loci that have so far been characterized are more often involved transcriptional regulation and cell cycle control (6,11,40). Consistent with this, some of our target genes are involved in modifying the epigenome. This includes *KANSL1* which acetylates lysine residues of nucleosomal histone H4. *KANSL1* has not been directly implicated in ovarian cancer biology previously, although it has been shown to form a fusion gene product with a G-protein coupled receptor for corticotropin-releasing factor, CRH1 in a small proportion of primary EOCs (41). In addition, *KANSL1* and other several of the candidate susceptibility genes are involved in mitosis and genome replication, biological processes known to play a key role in HGSOC development (42). Perturbations in vital cell cycle checkpoints allow cells to enter mitosis in spite of DNA damage, enabling cells to acquire additional genetic alterations that promote neoplastic development. In the splice-TWAS we identified *PRC1*, a key regulator of cytokinesis required for spindle formation which reportedly acts as an oncogene during bladder cancer development (43). Strikingly, in data from a knockout screen *PRC1* shows similar levels of essentiality as *MYC* (a known essential gene and likely GWAS target gene in HGSOC (44)) strongly indicating *PRC1* plays a functional role in the development of ovarian tumors. In this same functional category is *CHMP4C* (chromatin-modifying protein 4C), a protein involved in the final steps of cell division, coordinating midbody resolution with the abscission checkpoint (45). *CHMP4C* expression has been implicated in several cancers and has been proposed as a diagnostic tumor marker and therapeutic target for ovarian cancer (46,47).

In summary, we have performed a TWAS based on the integration of GWAS data for HGSOC and gene expression data for both normal and tumor tissues associated with HGSOC pathogenesis, to identify candidate susceptibility genes associated with HGSOC risk alleles. Uniquely, this study established splice-TWAS associations (in addition to total expression associations) as a major component of HGSOC heritability, highlighting the significance of a variety mechanisms of transcriptional regulation at GWAS risk loci. This study emphasizes the significance of tissue-specific transcript wide expression and the need to evaluate both normal and cancer tissues associated with disease pathogenesis to identify candidate susceptibility genes, principles that apply not just to the analysis of ovarian cancer, but to many other phenotypes.

## METHODS

### Data processing and QC

#### Genotypes

Germline DNA from normal OSEC and FTSEC samples were genotyped using the Oncoarray platform (10). For TCGA data, SNP genotype calls using Birdsuite were downloaded from the TCGA legacy archive and imputed using the EAGLE pipeline provided by the Michigan imputation server. The following genotype QC was performed across all studies: SNPs were retained if they had imputation INFO>0.9; locus missingness <5%; Hardy-Weinberg equilibrium p-value > 5×10^−6^; and minor allele frequency > 1% (thresholds based on GTEx Consortium recommendations). Individuals were excluded if they had more than 5% missing sites. Two genotype principal components were computed to account for ancestry and included as covariates in all subsequent analyses.

#### Gene/exon expression in HGSOC precursor tissues

OSECs and FTSECs were harvested from histologically normal ovaries and fallopian tubes removed from women diagnosed with ovarian, uterine or cervical cancer. Short-term cultures were established (48,49). OSECs were harvested using a cytobrush and cultured in NOSE-CM media containing 15% fetal bovine serum (FBS, Hyclone), 34 μg ml^−1^ bovine pituitary extract, 10 ng ml^−1^ epidermal growth factor (Life Technologies), 5 μg ml^−1^ insulin and 500 ng ml^−1^ hydrocortisone (Sigma-Aldrich). Fallopian epithelia were dissociated from stromal tissues by Pronase/DNase I digestion (Roche and Sigma-Aldrich, respectively) for 48-72 hours at 4˚C. Purified epithelia were cultured on collagen I (Sigma-Aldrich) using DMEM/F12 base media supplemented with 2% Ultroser G (Pall Corporation). At ∼80% confluency, cells were lysed using the QIAzol reagent and RNA extracted using the RNeasy Mini kit (both QIAgen). RNA sequencing was performed by the University of Southern California Epigenome Core Facility using 50bp single end reads. All data processing was performed using ‘R’ and ‘Bioconductor’, and packages therein.

RNAseq data for 394 HGSOC samples was obtained from The Cancer Genome Atlas (TCGA) data portal as protected data (raw sequencing, fastq files) and downloaded via CGHub’s geneTorrent. Data was aligned to a reference genome (hg19) using STAR. And quality control of aligned samples performed using RSeQC. GC bias and batch effect corrections were performed using EDASeq and ‘sva’. To adjust for batch effects we used an empirical Bayes framework (comBat), available in ‘sva’.

Exon-level events were called directly from the aligned reads using LeafCutter (50) with default parameters. LeafCutter was then used to compute intron excision ratios and remove any introns used in <40% of samples. Finally, all excision ratios were quantile normalized and three expression principal components were computed and included as covariates in all subsequent analyses to account for latent confounders.

#### Gene/exon expression in non-ovarian TCGA samples

Normalized gene and exon level events were downloaded from the TCGA FireCloud. Exon usage was previously quantified using MapSplice. Finally, all expression/exon measurements were quantile normalized. As with the ovarian data, all expression/exon measurements were quantile normalized and three expression principal components were computed and used as covariates in all subsequent analyses.

#### Gene expression in GTEx samples

Processed and normalized expression and genotypes were downloaded from dbGAP and the GTEx Portal as described in (51). For each tissue the following covariates were included in all analyses: three genetic principal components, sex, platform, and 14-35 expression factors (52) as selected by the main GTEx analysis.

### Heritability and genetic correlation

Mean heritability and genetic correlation were estimated using Haseman-Elston regression (53) as implemented in GCTA (54). All SNPs within 500kb of the gene boundary were used to define the cis locus and construct the corresponding kinship matrix. Standard errors for genetic correlation were estimated as in (55).

### TWAS analyses

#### GWAS data

GWAS data from the Ovarian Cancer Association Consortium as described in (10) was downloaded and aligned to hg19 HapMap3 SNPs (excluding A/T or C/G SNPs due to strand ambiguity). These SNPs are consistently imputed with high accuracy across diverse genotyping platforms and were used to compute all TWAS weights.

#### TWAS predictors

TWAS predictors were computed using the FUSION software (see Data Availability). Briefly, for each gene or splice variant, SNPs from +/−500kb of the feature boundary were extracted and used to estimated cis-SNP-heritability (54). Features that had nominally significant cis-SNP-heritability (LRT P<0.01) were retained, and the genotypes were used to train TWAS predictive models using BLUP, elastic net, BSLMM (for BRCA data only) and LASSO models. Five-fold cross-validation was performed for each model and the best scoring model (out-of-sample) was used for the TWAS test. Across all tissues and features, the top eQTL was the best predictor only 26% of the time. Surprisingly, the BLUP predictor - which has the weakest penalization in favor of sparsity - was the most common best predictor (best 33% of the time), suggesting a greater degree of effect heterogeneity in this data than studies of normals where cis-expression effects are typically sparse (56). For GTEx analyses, TWAS expression weights were previously computed as described in (13), and downloaded from the FUSION website.

#### TWAS tests

The FUSION software was used to perform TWAS tests across all predictive models (14). Models were considered “transcriptome-wide significant” if they passed Bonferroni correction for all 68,132 genes and exon events tested.

Summary-based conditional analyses between TWAS and GWAS associations were performed using FUSION (57). For a given significant TWAS association, the gene/exon expression was predicted into the 1000 Genomes EUR samples to estimate the LD between the predicted model and each SNP in the locus. Each GWAS SNP was then conditioned on the predicted model using the LD estimate to quantify the amount of residual association signal. Stepwise model selection was performed by including each TWAS-associated feature (from most significant to least) into the model until no feature remained conditionally significant.

Summary-based conditional analyses for individual SNPs were performed using GCTA-COJO (57). Colocalization analyses were performed using the COLOC software (58) and the marginal eQTL/spQTL statistics for a given feature.

### Functional analyses

Differential gene expression analyses were performed using the OSEC, FTSEC and HGSOC data sets described above. Processed CERES knockout data were downloaded (18) and data for the 13 HGSOC lines included in this study were used in our analyses.

### Ethics Approval

This study was performed with the approval of the Cedars-Sinai Medical Center Institutional Review Board.

### Data and Code Availability

Code and documentation for all methods has been made available on the TWAS/FUSION web-site (http://gusevlab.org/projects/fusion/). Trained TWAS models for all genes and splice variants and corresponding OvCa TWAS association statistics will be made available on the TWAS/FUSION web-site upon publication.

## ACKNOWLEDGEMENTS

This work was supported by an R21 award from the NIH (R21CA22007801). The results shown here are in part based upon data generated by the TCGA Research Network: http://cancergenome.nih.gov/. Some of the normal tissue specimens were collected as part of the USC Jean Richardson Gynecologic Tissue and Fluid Repository, which is supported by a grant from the USC Department of Obstetrics & Gynecology and the NCT Cancer Center Shared Grant award P30 CA014089 (to the Norris Comprehensive Cancer Center). K.L. is supported in part by a K99/R00 Pathway to Independence Award from the NIH (R00CA184415) and institutional support from the Samuel Oschin Comprehensive Cancer Institute at Cedars-Sinai Medical Center.H.N. and M.A.S.F. are supported by grants 2015/07925-5 and 2017/08211-1 from Sao Paulo Research Foundation (FAPESP). H.N. is also supported by an institutional grant (Henry Ford Hospital). This work was supported in part by the Ovarian Cancer Research Fund Alliance Program Project Development Grant (373356): Co-Evolution of Epithelial Ovarian Cancer and Tumor Stroma. Additional support for this work came from NIH/NCI grants 1R01CA211707 and 1R01CA207456 and OCRF award 258807.

## SUPPLEMENTARY TABLE LEGENDS

**Supplementary Table S1: Number of samples/features and mean heritability in each analyzed cohort.**

**Supplementary Table S2: Genetic correlation across HGSOC subtypes.**

**Supplementary Table S3: Prediction accuracy for significantly heritable genes in each cohort.**

**Supplementary Table S4: Correlation of genetic predictors across cohorts for shared significantly heritable genes.**

**Supplementary Table S5: Significant TWAS associations for total expression. Supplementary Table S6: TWAS Z-score across all tissues for each significant TWAS gene.**

**Supplementary Table S7: Significant splice-TWAS associations for total expression.**

**Supplementary Table S8: Splice-TWAS Chi-squared statistic across all tissues for each gene with a significant splicing association.**

**Supplementary Table S9: Number of gene/splicing predictors evaluated in 13 genome-wide significant regions.**

## SUPPLEMENTARY FIGURE LEGENDS

**Supplementary Figure S1. TWAS association Manhattan plots.** Plot of physical position (x-axis) and TWAS association p-value (y-axis) across each expression reference panel. Transcriptome-wide significant associations highlighted in red.

**Supplementary Figure S2. Relationship between tested and significant genes.** The number of TWAS significant genes (y-axis) is plotted as a as a function of the total number of significantly heritable (predictable) genes (x-axis). The black line indicates linear regression fit. SP: splicing; BRCA: breast; OVCA: ovarian; PRAD: prostate.

**Supplementary Figure S3: Conditional analysis of TWAS association for *RCCD1*.** (top) Local Manhattan plot for GWAS association before (gray) and after (blue) conditioning on the predicted *RCCD1* expression from GTEx. (bottom) corresponding local Manhattan plot for eQTL associations to *RCCD1*.

**Supplementary Figure S4: Conditional analysis of TWAS association for *DNAL1*.** (top) Local Manhattan plot for GWAS association before (gray) and after (blue) conditioning on the predicted *DNAL1* expression from GTEx. (bottom) corresponding local Manhattan plot for eQTL associations to *DNAL1*.

**Supplementary Figure S5: Conditional analysis of splice-TWAS association for *INTS1*.** (top) Local Manhattan plot for GWAS association before (gray) and after (blue) conditioning on the predicted *INTS1* splicing in breast tumor. (bottom) corresponding local Manhattan plot for sp-QTL associations for this *INTS1* splicing event.

**Supplementary Figure S6: Correlation of predictors. A, B,** at the 17q21.31 locus; **C,** at the 19p13 locus. Heatmaps of pairwise predictor correlation for all transcriptome-wide significant genes/exons.

**Supplementary Figure S7: Conditional analysis of TWAS and sp-TWAS associations at chr19p13 locus. A,** Model selection was performed to identify four jointly significant TWAS associations (see Methods) highlighted in green. Local Manhattan plot shows GWAS signal before (gray) and after (blue) joint conditioning on the predicted expression of these four genes. **B**, (top) Local Manhattan plot for GWAS association before (gray) and after (blue) conditioning on the predicted *BABAM1* splicing in breast tumor. (bottom) corresponding local Manhattan plot for sp-QTL associations for this *BABAM1* splicing event.

